# The Impact of Acute High-Intensity Activity on Perceptual Decision-Making Dynamics

**DOI:** 10.1101/2023.02.14.528466

**Authors:** Davranche Karen, Giraud Dorian, Hays Arnaud, Gajdos Preuss Thibault

## Abstract

This study investigates the acute impact of high-intensity activity on perceptual decision-making, using computational modeling to assess changes during and after physical activity. Participants performed a two-alternative forced choice perceptual decision-making task at rest (pre- and post-exercise) and during six of eight 5-minute cycling bouts (totaling 47 minutes) under dual-task condition, while maintaining an average intensity of 86 ± 7% of their maximum heart rate. Drift diffusion modeling was applied to accuracy and reaction time data to estimate changes in evidence accumulation (drift rate), decision threshold (boundary separation), and non-decision processes (*t_er_*). Results revealed improved post-exercise performance, characterized by shorter non-decision time, potentially reflecting a transient improvement in motor or perceptual efficiency. During ongoing physical activity, results indicate that exercise is associated with a decrease in non-decision time and an increase in the efficiency of evidence accumulation, while response caution remains stable. These findings provide novel insights into how sustained high-intensity exercise modulates perceptual decision-making dynamics under physiological stress.

## Introduction

April 15, 2019. Notre-Dame is ablaze. After a strenuous climb, during which a firefighter analyzes and interprets the situation, she finally reaches the top of the tower. Now, she must decide which side of the raging fire to attack first. In such extreme conditions, is her decision-making affected by sustained physical and mental demand? And if so, which specific processes are altered during and after such intense exercise? These questions motivate our investigation into how high-intensity exercise influences perceptual decision-making.

It is well established that acute moderate-intensity physical activity can enhance cognitive performance across various cognitive tasks, including memory, vigilance, and conflict resolution (e.g., McMorris, 2021; Pontifex et al., 2019). However, the effects of high-intensity aerobic activity remain less clear. While some studies report impairments, others suggest null effects or even improvements (McMorris, 2021; Moreau & Chou, 2019). Recent reviews (Sudo, Costello, McMorris & Ando, 2022; Zheng et al., 2021; Pontifex et al., 2019) emphasize the need for more research into this intensity range, as most existing findings focus on moderate (58% of the literature) intensities of physical activity. This emphasis, often driven by methodological concerns, has left a gap in our understanding of how demanding physical exertion affects cognition.

High-intensity activity, typically defined as physiological demand exceeding 80% maximal oxygen uptake or HRmax (McMorris et al., 2016), triggers more pronounced physiological responses than conventional aerobic activity (Herold et al., 2019). These responses include changes in circulating lactate, blood glucose, circulating cortisol levels, or neurotrophic factors (Hashimoto et al., 2021; Singh & Staines, 2015), which may differentially modulate cognitive function. Such changes can affect various neural processes, potentially enhancing sensory encoding, increasing arousal, and modulating motor readiness.

Because cognitive processes cannot be directly observed, we used a computational modeling approach to infer them from behavioral data. Specifically, we applied the Drift Diffusion Model (DDM, Ratcliff, 1978) a well-validated model for analyzing two-alternative forced-choice tasks (Ratcliff et al., 2016). The DDM assumes that decisions result from the accumulation of noisy evidence until a response threshold is reached (Bogacz et al., 2006). The basic DDM contains four key parameters, each related to a different cognitive process: (1) the drift rate (v), reflecting the efficiency of evidence accumulation; (2) the decision threshold (a), representing response caution; (3) the starting point (z), indicating decision bias; and (4) the non-decision time (*t_er_*), encompassing stimulus encoding and motor execution. Participants completed a random dot motion task before, during, and after engaging in 5-minute cycling bouts at >80% HRmax. Behavioral data (accuracy and reaction times, RT) were used to estimate model parameters and track changes in cognitive processes across conditions.

## Materials and methods

### Participants

Nineteen young adults (one woman, age range: 18–34; mean age: 25 years) voluntarily participated in the experiment. All participants were healthy and practiced regular physical activities (peak oxygen uptake: 55 ± 8 mL kg−1 min−1; maximal heart rate: 184 ± 9 bpm). All participants were fully informed about the study and signed a written informed consent before inclusion. The study was approved by the Ethics Committee for Research in Science and Technology of Physical Sports Activities (IRB00012476-2022-21-01-148).

### Random dot motion task (RDK)

We used a perceptual decision-making task, the random dot motion task (RDK), for which the DDM has consistently provided excellent accounts of behavioral data. The experiment was programmed in Python, utilizing components of the PsychoPy toolbox (Peirce et al., 2019), and was executed on a PC computer operating natively on Windows 10 with a 19.5-inch screen. Participant responses were recorded using two buttons (one for each hand).

The task requires participants to determine the global direction of a random dot kinematogram (RDK) featuring a proportion of dots moving coherently in the left or right signal direction. Each trial started with the presentation of the random dot motion stimulus, which remained on the screen until the participant responded. A trial is correct if the participant report the direction of coherently moving dots. A response time deadline was set to 5 s. The interval between the response to the stimulus and the next trial was 1.5 s. The coherence parameter (proportion of dots moving in the same direction) was set for each participant at the beginning of the experiment according to a 2 up and 1 down staircase method lasting 30 trials, which corresponds to an accuracy close to 70%.This participant-specific adjustment of task difficulty helps eliminate potential confounding factors arising from inter-individual differences in task performance, thereby making it easier to interpret the model parameters—and, in turn, the effects of the time window on these parameters. White dots were presented within a virtual 12.6° circular aperture centered on a 24.8° 3 13.9° black field. Each dot was a 4 x 4 pixel (0.05° square), moving at a speed of 8°/s. Dot density was fixed at 16.7 dots/deg2/s. Random dot motion was controlled by a white noise algorithm (Pilly & Seitz, 2009).

### Procedure

Participants made three separate visits to the laboratory, each spaced at least 48 hours apart: a characterization session, a familiarization session, and an experimental session (figure 1). Participants were positioned on an upright cycle ergometer (Lode Excalibur Sport, Groningen, The Netherlands), equipped with two thumb response buttons on the right and left handle grips. A computer screen was positioned at eye level, approximately 1 meter in front of the participant.

**Figure 1.**
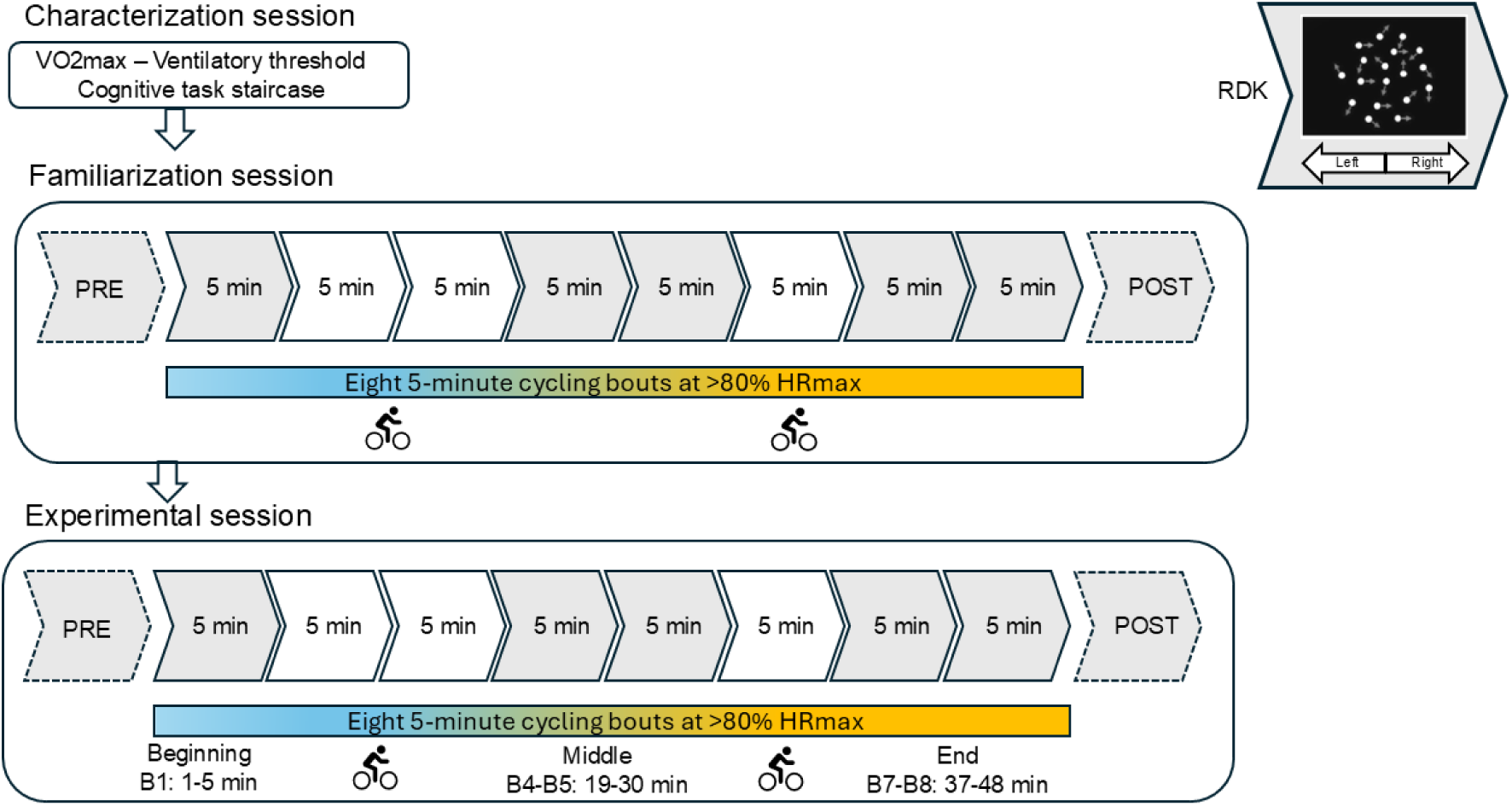
Overview of the experimental design and task structure. The study included three laboratory visits: (1) a characterization session to assess individual physiological parameters and calibrate the difficulty of the cognitive task, (2) a familiarization session, identical in all respects to the experimental session, aimed at stabilizing performance, and (3) an experimental session combining 5-minute cycling bouts at >80% HRmax with (in grey) or without (in white) cognitive testing. Cognitive performance was assessed at rest for 5 minutes before (PRE) and immediately after exercise (POST), and during exercise at three-time windows: Beginning (minutes 1–5, B1), Middle (minutes 19–30, B4–B5), and End (minutes 37–48, B7–B8) of the physical exercise. A schematic representation of the random dot motion task (RDK) is shown in the top right corner of the figure.

The preliminary session was conducted to individually adjust the difficulty of the cognitive task, assess V̇O2max, and determine ventilatory thresholds (Wasserman, 2012) (see table 1 for details). Participants were seated on the cycle ergometer, fitted with a facemask to measure gas exchange data (Innocor CO, Cosmed, Italia) and a heart rate monitor. The incremental cycling activity to exhaustion test started at an initial power level defined according to the participant’s level (100 watts for non-cyclists and 150 watts for cyclists) and increased by 30 watts every 2 minutes until exhaustion. The end of the test was determined by volitional cessation of exercise or failure to maintain pedal cadence above 60 rpm despite strong verbal encouragement. During the first step of this maximal test, 30 trials of the RDK task were performed to set up the consistency parameter according to a staircase procedure.

**Table 1.**
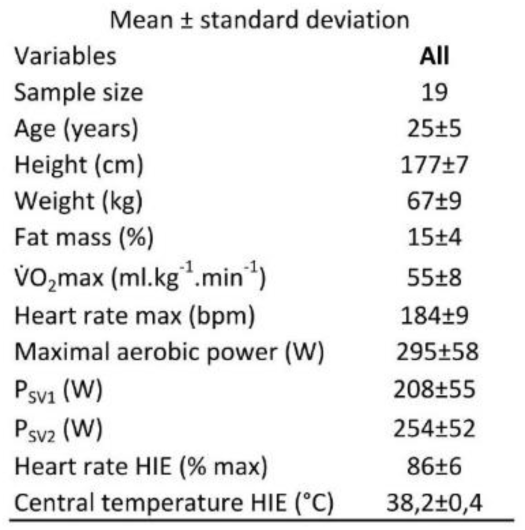
Anthropometric and physiological characteristics of participants. Note. P_SV_: Power corresponding to the first SV1 and second SV2 ventilatory thresholds, HIE: high-intensity exercise, V̇O2max: maximal oxygen consumption.

Before the experimental session, a familiarization session (about 1,250 trials), identical in all respects to the experimental session, was conducted to stabilize performance and control for potential learning effects, as well as to assess the physiological load of the dual task performed while cycling. The data of the familiarization session were not included in the analyses.

The experimental protocol involved high-intensity interval activity, a discontinuous mode of endurance exercise characterized by short bouts of high-intensity workloads interspersed with periods of rest. Participants completed a perceptual decision-making task at rest both before and after eight 5-minute bouts of cycling. Each 5-minute cycling bout was followed by a 1-minute rest period, resulting in a total session duration of 47 minutes. During six of the eight cycling bouts, participants performed the perceptual decision-making task while cycling at a workload fixed at 90% of SV1—an intensity determined during the incremental test of the preliminary session—placing them in a dual-task situation involving both cognitive and physical activity. Cardiorespiratory parameters were continuously recorded, and capillary blood samples from the fingertip were collected before and immediately after each of the 5-minute cycling bouts for the measurement of lactate and glucose levels. The skin was prepared by cleaning with alcohol, allowed to dry, and then punctured with an automated lancet. Blood samples were analyzed using a Biosen blood analyzer (EKF Diagnostics, UK).

To examine the time-dependent effects of exercise on cognitive processes, the dual-task period was segmented into three time windows: the Beginning phase (B1), corresponding to the first 5 minutes of the physical activity; the Middle phase, spanning minutes 19 to 30 (B4-B5); and the End phase, encompassing minutes 37 to 48 (B7-B8). This segmentation was based on the expected temporal dynamics of physiological responses during sustained intermittent cycling. The Beginning of exercise (B1) was chosen as a baseline, as it corresponds to the early phase of exertion when physiological changes (e.g., blood lactate accumulation, cardiovascular strain) have not yet fully developed, allowing us to capture the initial dual-task condition under minimal metabolic load. The Middle window reflects a phase where physiological responses typically plateau (e.g., elevated but stabilized lactate and heart rate), while the End window captures potential cumulative effects of prolonged exertion and fatigue. This time window approach enables us to better track the evolving impact of physical exercise on decision-making processes over time.

### Data Analyses

The experiment was designed to evaluate the exercise-induced effects on cognitive processes throughout high-intensity activity and immediately afterward exercise bout. Due to the interplay between physical demands and cognitive performance in the dual-task condition, cognitive data recorded at rest (pre- and post-exercise, 4916 trials) and during the dual-task condition (blocks while cycling, 12402 trials) were analyzed separately. During the physical activity, we anticipated that the impacts on decision-making processes may vary according to time spent on task. To capture the time-specific effects of exercise on cognitive performance, we conducted analyses across three specific time windows.

Anticipations (RTs < 200 ms; 0.13 %) were discarded from all analyses. Five subjects did not perform the perceptual task appropriately (i.e., exhibiting a percentage of accuracy below 50% or employing a stereotyped response strategy). They were excluded from the initial sample of 24 participants. Regarding physiological measures, we analyzed glucose and lactate levels within linear mixed-effects models. These models included the ‘sample’ (PRE and B1 to B8) expressed in mmol/L as a fixed effect, with ‘subject’ as random effect.

The parameters of the Drift Diffusion Model (DDM) were estimated using a hierarchical Bayesian approach implemented with the HSSM toolbox in Python (Fengler et al., 2025). Two separate sets of model fits were conducted: one comparing the PRE and POST time windows, and another comparing the Beginning, Middle, and End phases of exercise. In both cases, the fitting procedure was identical. We evaluated eight versions of a DDM without inter-trial variability. The first was a “basic” model, in which all parameters—starting point (*z*), drift rate (*v*), threshold (*a*), and non-decision time (*t_er_*)—were held constant across blocks. The “full” model allows all parameters, except the starting point, to vary across time windows. The remaining six models each allowed different combinations of these parameters to vary across time windows. For example, the “*vt*” model permits both the drift rate (*v*) and non-decision time (*t_er_*) to vary across time windows, and is formally defined in HSSM as:

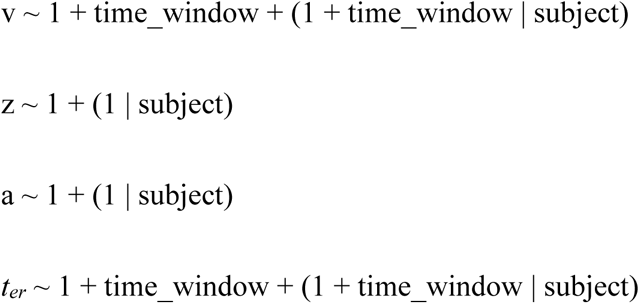

For all models, we used the default priors provided by HSSM. We then assessed convergence by visually inspecting the Markov Chain Monte Carlo (MCMC) traces and by examining the R-hat statistic from Brooks and Gelman (1998). Model comparison was carried out using the Bayesian Information Criterion (BIC). Subsequently, we evaluated model fits by visually comparing the best-fitting model to the basic model and analyzed the parameter estimates of the best model. This procedure provided two complementary approaches to assess the influence of experimental blocks on the parameters: (i) identifying which model structure best captures the data and (ii) examining the extent to which parameter estimates depend on time windows.

## Results

### Physiological measures

#### Heart rate

The linear mixed model revealed a significant effect of block on heart rate (F(7, 133) = 16.83, p < .001). Post-hoc comparisons showed that heart rate at block B1 was significantly lower than at all subsequent blocks (B3 to B8; all p < 0.001), while the difference between B1 and B2 was not significant (p = 0.44). Heart rate at B2 was also significantly lower than at blocks B4 to B8 (p < 0.05), but not significantly different from B3 (p = 0.21; see table 2 for means and standard deviation). No significant differences were found between any other block pairs from B3 to B8, suggesting that heart rate stabilized after the second or third block (see appendix S1 for individual trajectories across the protocol).

**Table 2.**
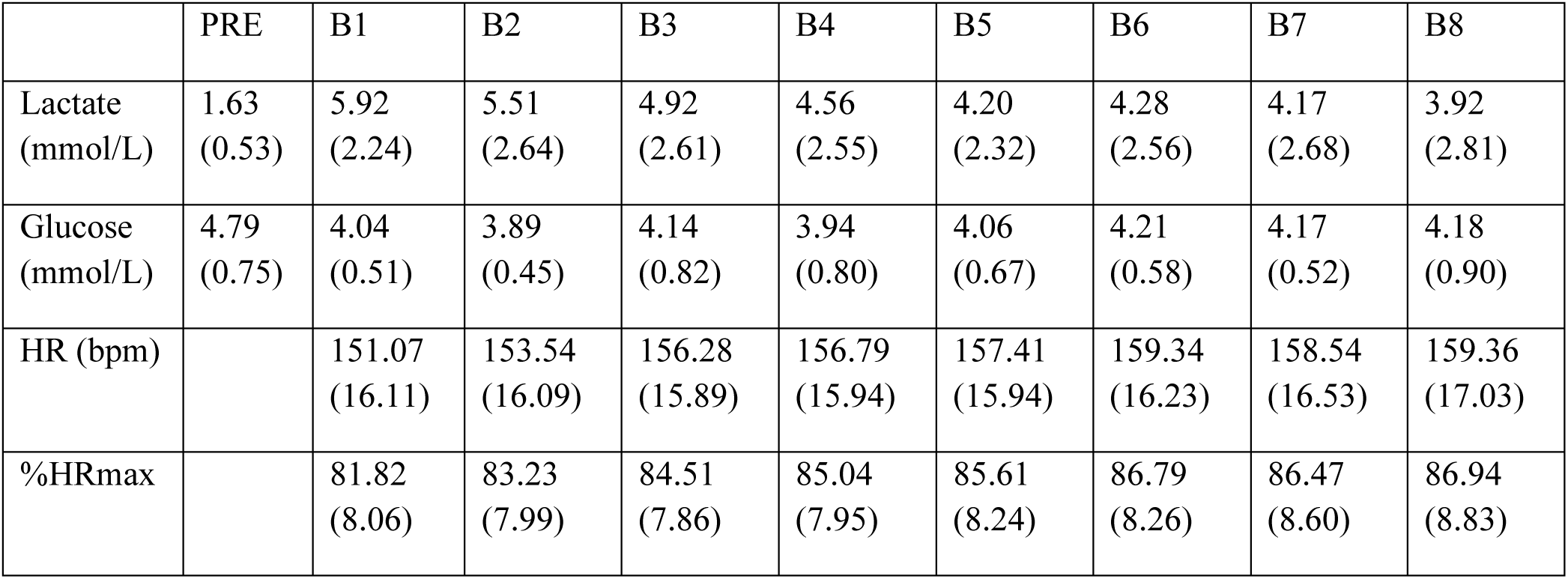
Mean values of glucose, lactate, heart rate (HR), and percentage of maximum heart rate (%HRmax) across bouts of high-intensity exercise. Note. Values represent the mean (standard deviation) for each bout.

|  | PRE | B1 | B2 | B3 | B4 | B5 | B6 | B7 | B8 |
| --- | --- | --- | --- | --- | --- | --- | --- | --- | --- |
| Lactate (mmol/L) | 1.63<br>(0.53) | 5.92<br>(2.24) | 5.51<br>(2.64) | 4.92<br>(2.61) | 4.56<br>(2.55) | 4.20<br>(2.32) | 4.28<br>(2.56) | 4.17<br>(2.68) | 3.92<br>(2.81) |
| Glucose (mmol/L) | 4.79<br>(0.75) | 4.04<br>(0.51) | 3.89<br>(0.45) | 4.14<br>(0.82) | 3.94<br>(0.80) | 4.06<br>(0.67) | 4.21<br>(0.58) | 4.17<br>(0.52) | 4.18<br>(0.90) |
| HR (bpm) |  | 151.07<br>(16.11) | 153.54<br>(16.09) | 156.28<br>(15.89) | 156.79<br>(15.94) | 157.41<br>(15.94) | 159.34<br>(16.23) | 158.54<br>(16.53) | 159.36<br>(17.03) |
| %HRmax |  | 81.82<br>(8.06) | 83.23<br>(7.99) | 84.51<br>(7.86) | 85.04<br>(7.95) | 85.61<br>(8.24) | 86.79<br>(8.26) | 86.47<br>(8.60) | 86.94<br>(8.83) |

#### Blood Biomarkers

Linear mixed-effects models revealed a significant effect of exercise on both blood glucose (F(8, 127) = 5.76, p < 0.001) and blood lactate levels (F(8, 127) = 5.76, p < 0.001), measured at the end of each 5-minute cycling bout (table 2 and Supplementary Figures S1–S3). Post-hoc comparisons showed that glucose concentrations were significantly higher at the pre-exercise baseline compared to all subsequent time points (B1 to B8; all p < 0.01), after which values remained relatively stable. In contrast, lactate concentrations were significantly lower at the pre-exercise baseline than at all subsequent time points (B1 to B8; all p < 0.01), indicating a sharp increase after exercise onset followed by a plateau.

### Computational modeling results

#### Pre- and Post-exercise Effects

BIC model comparison shows that models that allow non-decision time (*t_er_*) to vary between PRE and POST time windows clearly dominate those who don’t (table 3). There are, however, little differences between the models that allow *t_er_* to vary across time windows, which suggest that threshold and drift variations do not contribute to improve fits.

**Table 3.** Bayesian Information Criterion (BIC) for each model. In the “basic” model, all parameters—starting point (z), drift rate (v), threshold (a), and non-decision time (t_er_)—were held constant across PRE and POST time windows. The “full” model allows all parameters, except the starting point, to vary across time windows. The remaining six models each allowed different combinations of these parameters to vary across time windows. For example, the “vt” model permits both the drift rate (v) and non-decision time (t_er_) to vary across time windows.

| Model | BIC |
| --- | --- |
| at | 3585 |
| vt | 3590 |
| full | 3593 |
| $t_{er}$ | 3602 |
| a | 3730 |
| av | 3733 |
| v | 3932 |
| basic | 4044 |

However, quantile probability plots suggest that letting the threshold *a* and the non-decision time *t_er_* vary across time windows does not dramatically improve the fit quality compared to the basic model (Figure 2 and Supplementary Figure S6).

**Figure 2.**
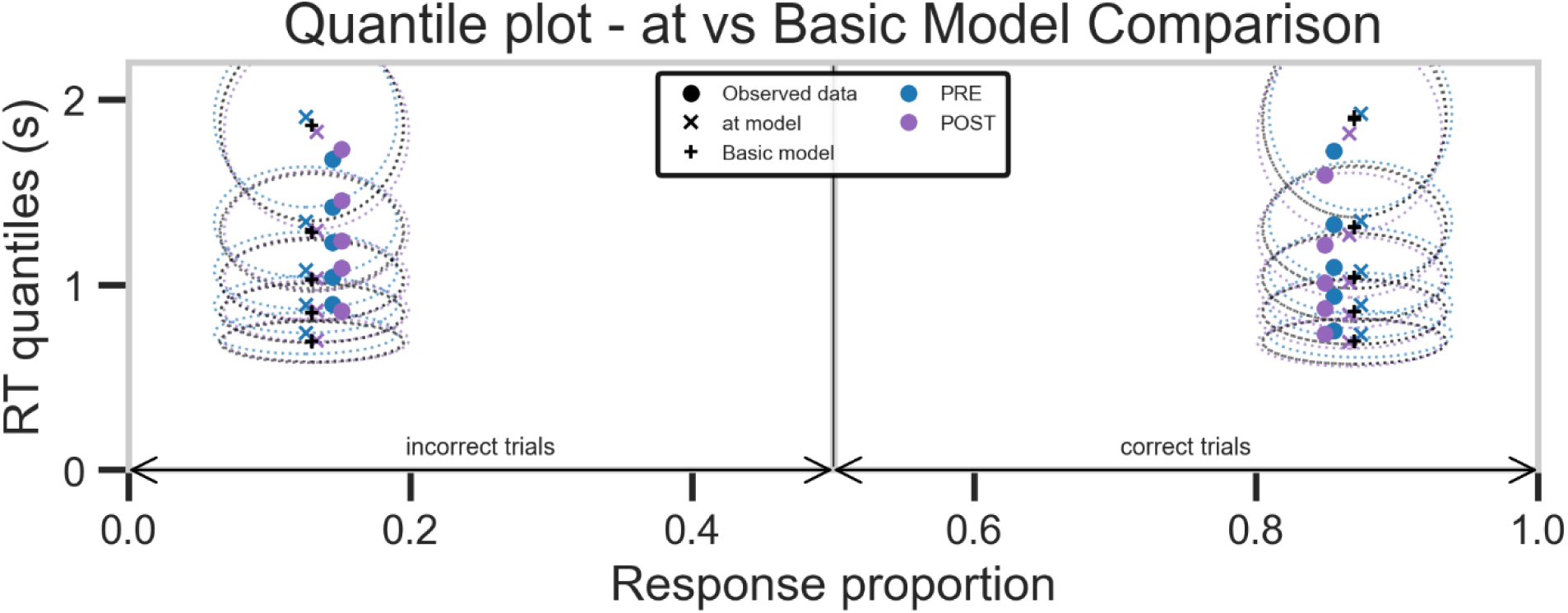
Quantile probability functions averaged across subjects for before (PRE, in blue) and after (POST, in purple) exercise, constructed by plotting RT quantiles (.1, .3, .5, .7, .9; y-axis) of the distributions of correct and incorrect responses against the corresponding response type proportion (x-axis). Dots represent observed data, “+” represent predictions of the basic model, and “x” represent predictions of the at model (using in both cases best fitting parameters). Dotted ellipses around the crosses represent 95% confidence intervals, reflecting the expected variability arising from model stochasticity and parameter estimation uncertainty.

The results of the fit of the best model (*at* model) confirm this assessment (figure 3; see Table S2 in appendix for a full summary of the results). We found that 82% of the posterior for the boundary parameter (*a*) difference between POST and PRE time windows was below 0, with a very small mean effect (M = -0.033, 94% HDI: [−0.099, 0.42]) compared to the average value across time windows (M = 0.979, 94% HDI: [0.839, 1.114]). While the effect on the non-decision time (*t_er_*) is more significant, with 94% of the posterior between POST and PRE time windows below 0, its amplitude is small (M = -0.032, 94% HDI: [−0.069, 0.008]) compared to the average value across time windows (M = 0.50, 94% HDI: [0.43, 0.58]).

**Figure 3.**
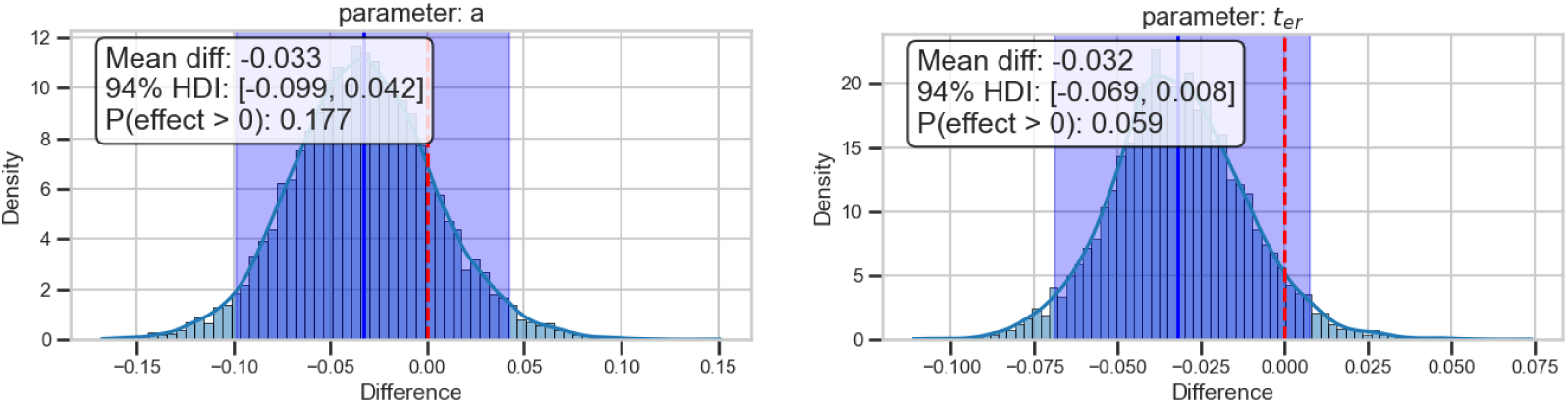
Posteriors for the boundary (a, left panel) and non-decision time (t_er_, right panel) parameters difference between POST and PRE time windows.

#### During-effort Effects

BIC model comparison shows that the model that allow drift rate (*v*) and non-decision time (*t_er_*) to vary across time windows is clearly the better model among those we compared (table 4).

**Table 4.** Bayesian Information Criterion (BIC) for each model. In the “basic” model, all parameters—starting point (z), drift rate (v), threshold (a), and non-decision time (t_er_)—were held constant across time windows. The “full” model allows all parameters, except the starting point, to vary across time windows. The remaining six models each allowed different combinations of these parameters to vary across time windows. For example, the “vt” model permits both the drift rate (v) and non-decision time (t_er_) to vary across time windows.

| Model | BIC |
| --- | --- |
| vt | 9655 |
| full | 9682 |
| at | 9710 |
| $t_{er}$ | 9721 |
| av | 9824 |
| a | 9851 |
| v | 9970 |
| basic | 10182 |

Visual inspection of quantile probability plots confirm that letting the non-decision time (*t_er_*) and drift rate (*v*) vary across time windows improve the fit quality (figure 4 and Supplementary Figure S7).

**Figure 4.**
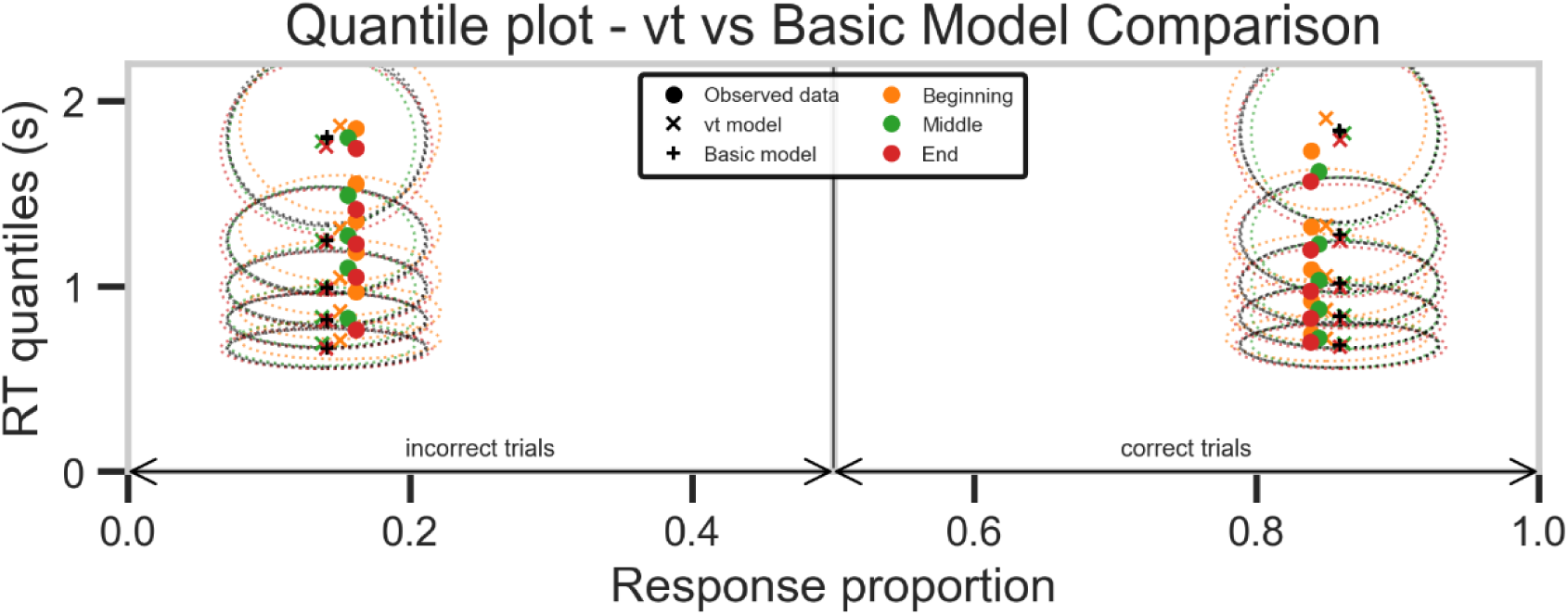
Quantile probability functions averaged across subjects at the Beginning (in orange), Middle (in green) and End (in red) of exercise, constructed by plotting RT quantiles (.1, .3, .5, .7,.9; y-axis) of the distributions of correct and incorrect responses against the corresponding response type proportion (x-axis). Dots represent observed data, “+” represent predictions of the basic model, and “x” represent predictions of the vt model (using in both cases best fitting parameters). Dotted ellipses around the crosses represent 95% confidence intervals, reflecting the expected variability arising from model stochasticity and parameter estimation uncertainty.

The results of the fit of the best model (*vt* model) confirm this assessment (figure 5; see Table S3 in appendix for a full summary of the results). We found that the drift rate (*v*) likely increases from the Beginning to the Middle time windows (94% of the posterior for the difference above 0; figure 5, left-top), with a size effect about 7% (M = 0.097, 94% HDI : [−0.017, 0.215]) compared to the average value across time windows (M = 1.427, 94% HDI : [1.037, 1.861]). There is, however, no evidence for an effect between the Middle and End time windows (61% of the posterior for the difference above 0; figure 5, left-middle). The non-decision time (*t_er_*) likely decreases between the Beginning and the End of exercise (97.5% of the posterior for the difference below 0; figure 5, right-bottom), with a size effect about 8% (M = - 0.041, 94% HDI : [−0.070, -0.004]) compared to the average value across time windows (M = 0.492, 94% HDI : [0.421, 0.563]). This result suggests an acceleration of perceptual-motor processes or motor facilitation induced by physical exertion. No clear difference was observed between the beginning and the Middle of exercise (81% of the posterior for the difference below 0; figure 5, right-top), nor between the Middle and end of exercise (73% of the posterior for the difference below 0; figure 5, right-middle), suggesting a stabilization of this effect.

**Figure 5.**
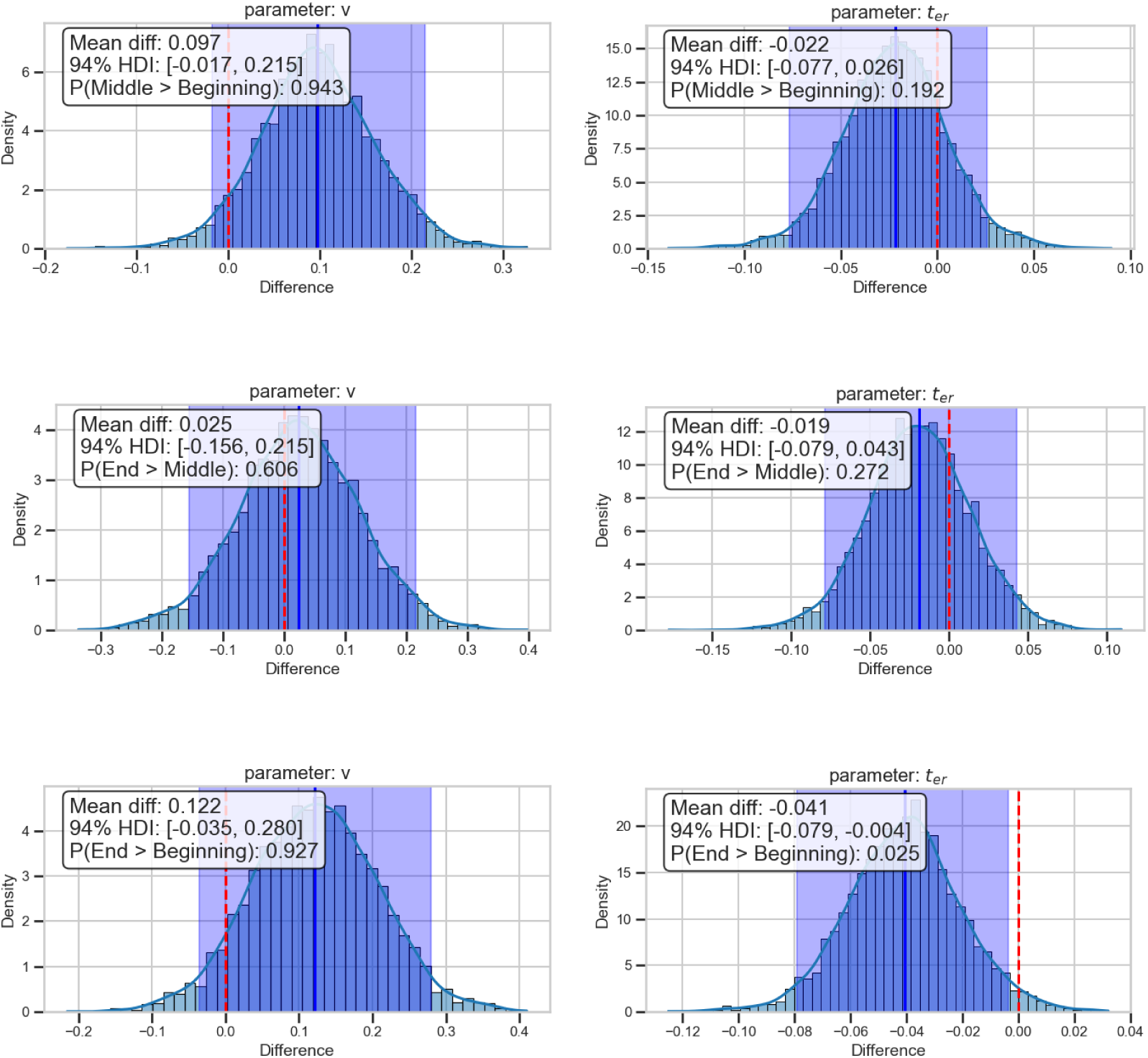
Posteriors for the drift rate (v, left panel) and non-decision time (t_er_, right panel) parameters difference across time windows.

## Discussion

### Summary

The present study aimed to investigate the impact of high-intensity activity on cognitive processes involved in perceptual decision-making. To this end, participants completed a perceptual decision task at rest (pre- and post-exercise), as well as six times during exercise in a dual-task condition involving both cognitive and physical activity. Cognitive processes were assessed by fitting a Drift Diffusion Model (DDM) to behavioral data collected across these time points. The transition from pre- to post-exercise conditions was associated with a possible reduction in non-decision time (*t_er_*), potentially reflecting a transient improvement in motor or perceptual efficiency. During ongoing physical activity, results indicate a decrease in non-decision time (*t_er_*) and an increase in the efficiency of evidence accumulation (drift rate, *v*), while response caution (boundary separation) remains stable.

### Post-effort modulation of cognitive processes

Findings indicate that transitioning from the Pre- to Post-exercise condition does not induce robust changes in decision-making parameters modeled by the DDM. However, a possible reduction in non-decision time (*t_er_*) was observed, which may reflect transient enhancements in sensory or motor efficiency. Bayesian model comparisons support a potential effect of exercise on non-decision components, with no consistent change core decision parameters. This suggests that exercise does not significantly impact perceptual discrimination processes, as measured by drift rate (*v*) or boundary separation (*a*).

This interpretation is consistent with prior evidence that cognitive benefits can persist briefly after the cessation of intense exercise (Tsukamoto et al., 2016a, 2016b; Moreau & Chou, 2019; McMorris et al., 2016), potentially due to sustained neurochemical or metabolic effects. Notably, elevated blood lactate has been shown to enhance motor system reactivity and sensory perception (Coco et al., 2009; 2016), possibly contributing to reduced *t_er_*. Increased cerebral blood flow and oxygenation following exercise may also support cognitive benefits (Rooks et al., 2010). Furthermore, catecholamine metabolism and the neuromodulatory role of lactate— produced even under fully aerobic conditions—warrant further exploration as potential mediators of the post-effort cognitive modulation.

### Ongoing physical activity impact on cognitive processes

Bayesian model comparisons suggest that sustained exercise is associated with a decrease in non-decision time (*t_er_*) and an increase in the efficiency of evidence accumulation (drift rate, *v*), while response caution (boundary separation, *a*) remains stable.

Drift rate analyses revealed a progressive increase in evidence accumulation from the Beginning to the Middle phase of exercise (minutes 19 to 30), followed by a plateau during the End phase (minutes 37 to 48), indicating changes in perceptual discrimination across exercise duration. In parallel, *t_er_* decreased from the beginning to the end of exercise, particularly between the early and middle phases, indicating a facilitation of perceptual-motor processes. The initial elevation in *t_er_* may reflect the time requiered for physiological adaptations, while the later stabilization could result from improved lactate clearance and a metabolic shift from carbohydrate to fat oxidation (Herold et al., 2022; Hashimoto et al., 2018; Tsukamoto et al., 2016b).

These results partially align with previous research demonstrating the influence of physiological states on cognitive functioning. For instance, Ratcliff & Van Dongen (2011), showed that reduced vigilance state induced by sleep deprivation accumulation via a reduction in *v*, while Du Rietz et al. (2019) reported a positive association between high-intensity activity and brain indices associated with arousal and attentional allocation. In contrast, the current findings suggest that exercise-induced changes are more nuanced and primarily reflected in non-decision processes.

This interpretation is consistent with prior evidence that physical exercise modulates both early (sensory) and late (motor) peripheral processing stages. Early studies have shown that moderate to intense exercise enhances sensory sensitivity and motor output speed, likely through increased excitability of the central nervous system (Davranche et al., 2005, 2006; Audiffren et al., 2008; Beyer et al., 2017). Exercise has been shown to increase cortical arousal and improve the signal-to-noise ratio in sensory processing, possibly due to the transient release of catecholamines and other neuromodulators (Davranche & Audiffren, 2004; Rodriguez et al., 2017; Moxon et al., 2007). At the motor level, evidence from single-pulse TMS studies suggests that exercise increases corticospinal excitability by reducing intracortical inhibition (Davranche et al., 2015), which could support more efficient motor command and response execution. These neurophysiological adaptations may underlie the observed improvements in *t_er_* and *v*, by rendering the brain more receptive to sensory input and more effective in translating decisions into action. Although the exact mechanisms remain to be fully elucidated, transient factors such as the release of catecholamines, serotonin, cortisol, or lactate production during exercise are likely contributors to these adaptive changes (Singh & Staines, 2015). Supporting this view, Coco et al. (2010) demonstrated that excitability of the primary motor cortex (M1), assessed via TMS over the hand area, was significantly enhanced at the end of a maximal cycling test, and that this enhancement strongly correlated with the increase in blood lactate levels. Interestingly, a similar effect on M1 excitability was also observed when lactate was increased via intravenous infusion in resting participants, suggesting a direct modulatory role of lactate on cortical excitability.

### Limitations and Perspectives

Several limitations warrant consideration. First, the sample was predominantly male (∼95%), which may limit the generalization of the findings to female populations. Nonetheless, previous studies involving gender-balanced cohorts have not reported sex-related differences in cognitive responses to elevated lactate levels (Perciavalle et al., 2015; Coco et al., 2016).

Second, there was substantial inter-individual variability in physiological responses to exercise at 90% of the first ventilatory threshold. Heart rates values ranged from 71% to 99% of HRmax, indicating variability in the relative contributions of aerobic and anaerobic metabolism. Post-hoc examination of individual lactate levels confirmed this variability, as participants were nearly evenly split between those above and below the 4 mmol/L threshold, commonly associated with the onset of blood lactate accumulation (OBLA). Since OBLA marks a key transition in metabolic pathways, this variability in lactate response could account for the absence of pre- vs post-exercise effect on *v* and may also obscure the effects during the different phases of the exercise. Supporting this interpretation, Coco et al. (2009) and Perciavalle et al (2014) reported that a large increase in blood lactate was associated with a significant worsening of attentional processes, suggesting that beyond a certain threshold—possibly near OBLA—rising lactate concentrations may no longer facilitate, but rather impair, cognitive functioning. This limit underscores the need for careful consideration of individual physiological profiles in future studies.

Additionally, differences in aerobic fitness likely modulate cognitive responses to intense exercise. Individuals with higher fitness levels exhibit more efficient lactate clearance, which may influence the duration and magnitude of post-exercise cognitive effects. Variability in tolerance to elevated lactate concentrations may also modulate cognitive outcomes, underscoring the importance of incorporating fitness assessments and lactate sensitivity measures in future research.

## Conclusion

By applying DDM to a perceptual decision-making task, this study provides new insights into the cognitive mechanisms underlying high-intensity exercise-induced changes. Our findings suggest that sustained physical activity is associated with changes in decision efficiency, primarily through a decrease in non-decision time and an increase in evidence accumulation. As a time-efficient and ecologically valid intervention, high-intensity exercise may represent a promising tool to transiently enhance cognitive function, warranting further investigation.

## Acknowledgements

We would like to thank the Filière Fastspor’in for its support in carrying out this study, and Anita Vergnaud et Laurine Stefanuto for their assistance in data collection.

## Declarations

### Funding

Not applicable.

### Conflicts of interest/Competing interests

Not applicable.

### Ethics approval

Ethics Committee for Research in Science and Technology of Physical Sports Activities (IRB00012476-2022-21-01-148).

### Consent to participate

All participants were fully informed about the study and signed a written informed consent before inclusion.

### Consent for publication

All authors have reviewed and approved the final manuscript for publication.

### Availability of data and materials

All data and scripts used in the present study are available at the following link: https://zenodo.org/records/17447883

### Code availability

Not applicable.

### Authors’ contributions

KD and TG contributed to conceptualization, methodology, data curation, formal analysis, and writing – original draft, review & editing. DG contributed to investigation. AH contributed to resources and investigation.

## Appendices

**Figure S1.**
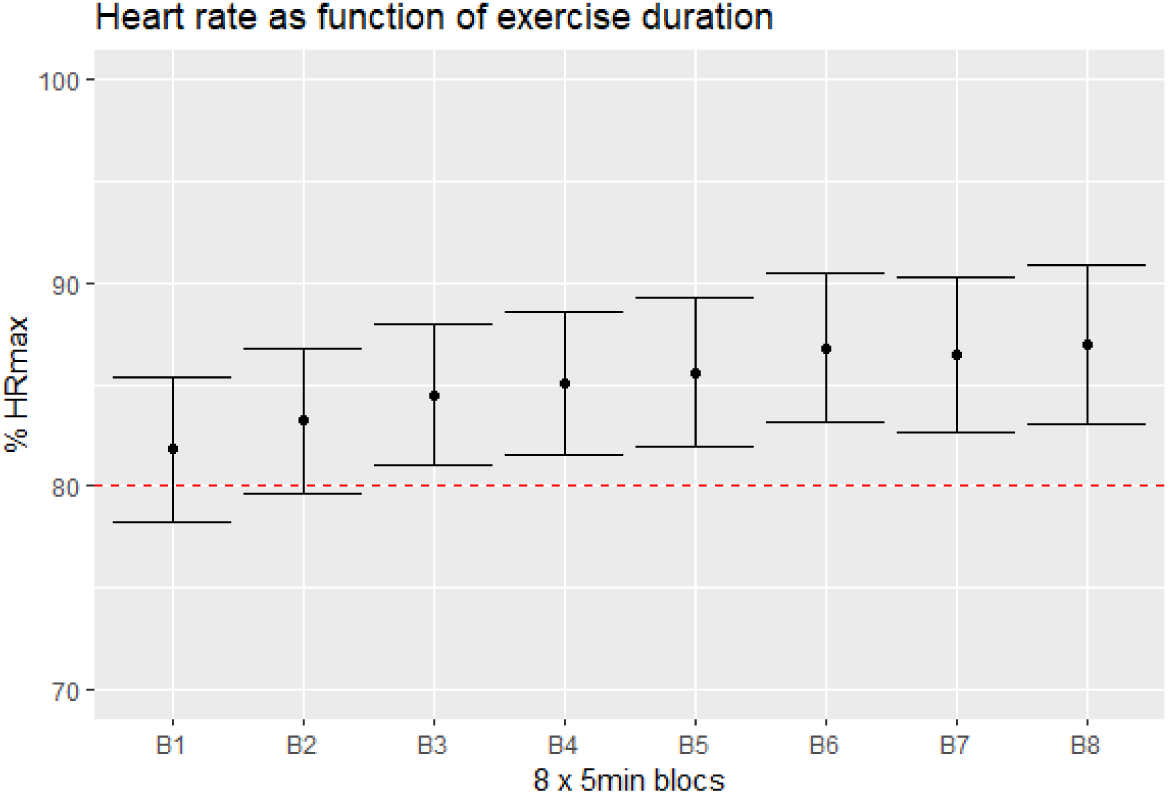
Heart rate (HR) across exercise blocks. Group-level mean (MD) ± standard deviation (SD).

**Figure S2.**
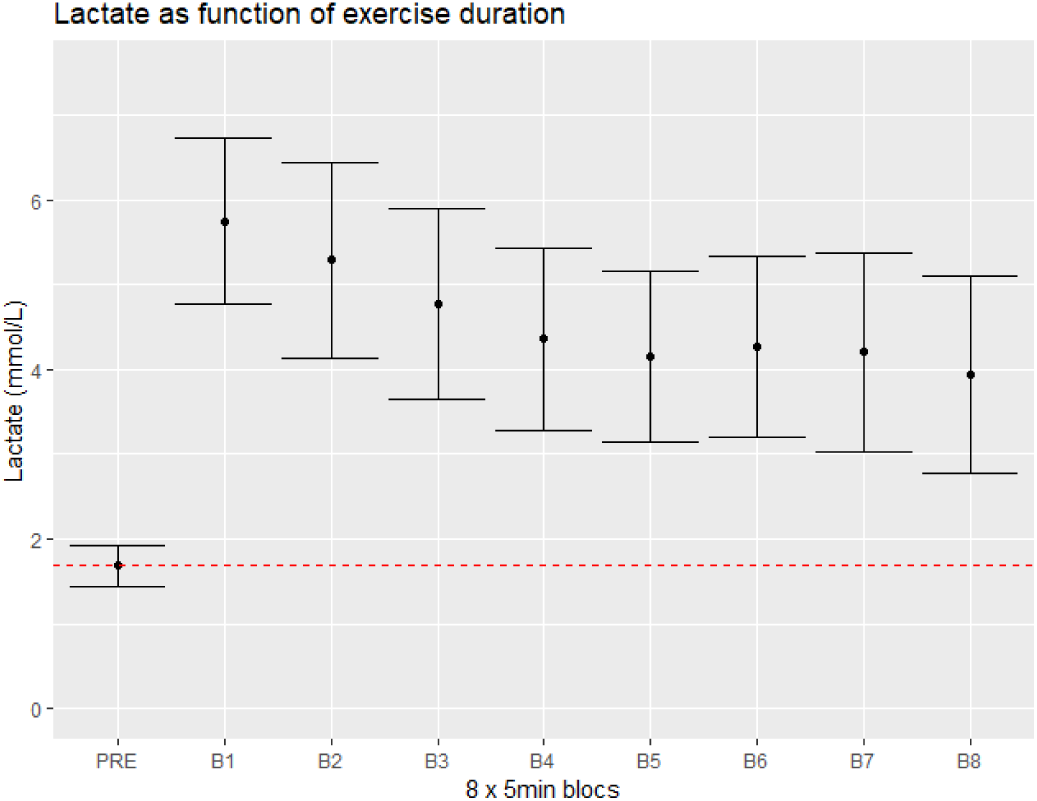
Blood lactate concentration across exercise blocks. Group-level mean (MD) ± standard deviation (SD).

**Table S2.** Full summary of the results of the fit of the best model (*at* model)

|  | mean | sd | hdi_3% | hdi_97% | ess_bulk | ess_tail | r_hat |
| --- | --- | --- | --- | --- | --- | --- | --- |
| **ter_C(condition)[POST]** | -0.032 | 0.020 | -0.069 | 0.008 | 2001.0 | 2991.0 | 1.0 |
| **z_Intercept** | 0.513 | 0.017 | 0.481 | 0.546 | 3476.0 | 4558.0 | 1.0 |
| **a_Intercept** | 0.979 | 0.073 | 0.839 | 1.114 | 1415.0 | 2595.0 | 1.0 |
| **a_C(condition)[POST]** | -0.033 | 0.037 | -0.099 | 0.042 | 3445.0 | 4855.0 | 1.0 |
| **t_Intercept** | 0.502 | 0.040 | 0.430 | 0.580 | 1419.0 | 2594.0 | 1.0 |
| **v_Intercept** | 1.500 | 0.214 | 1.108 | 1.903 | 1318.0 | 2601.0 | 1.0 |

**Figure S3.**
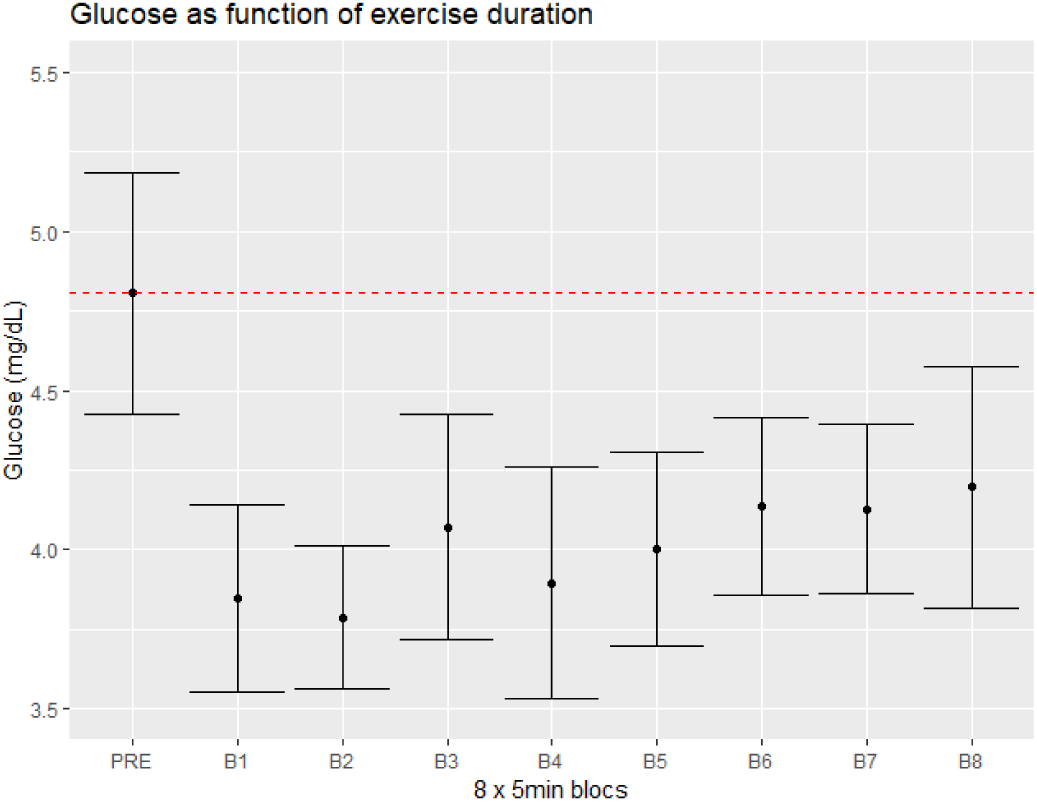
Blood glucose concentration across exercise blocks. Group-level mean (MD) ± standard deviation (SD).

**Figure S4.**
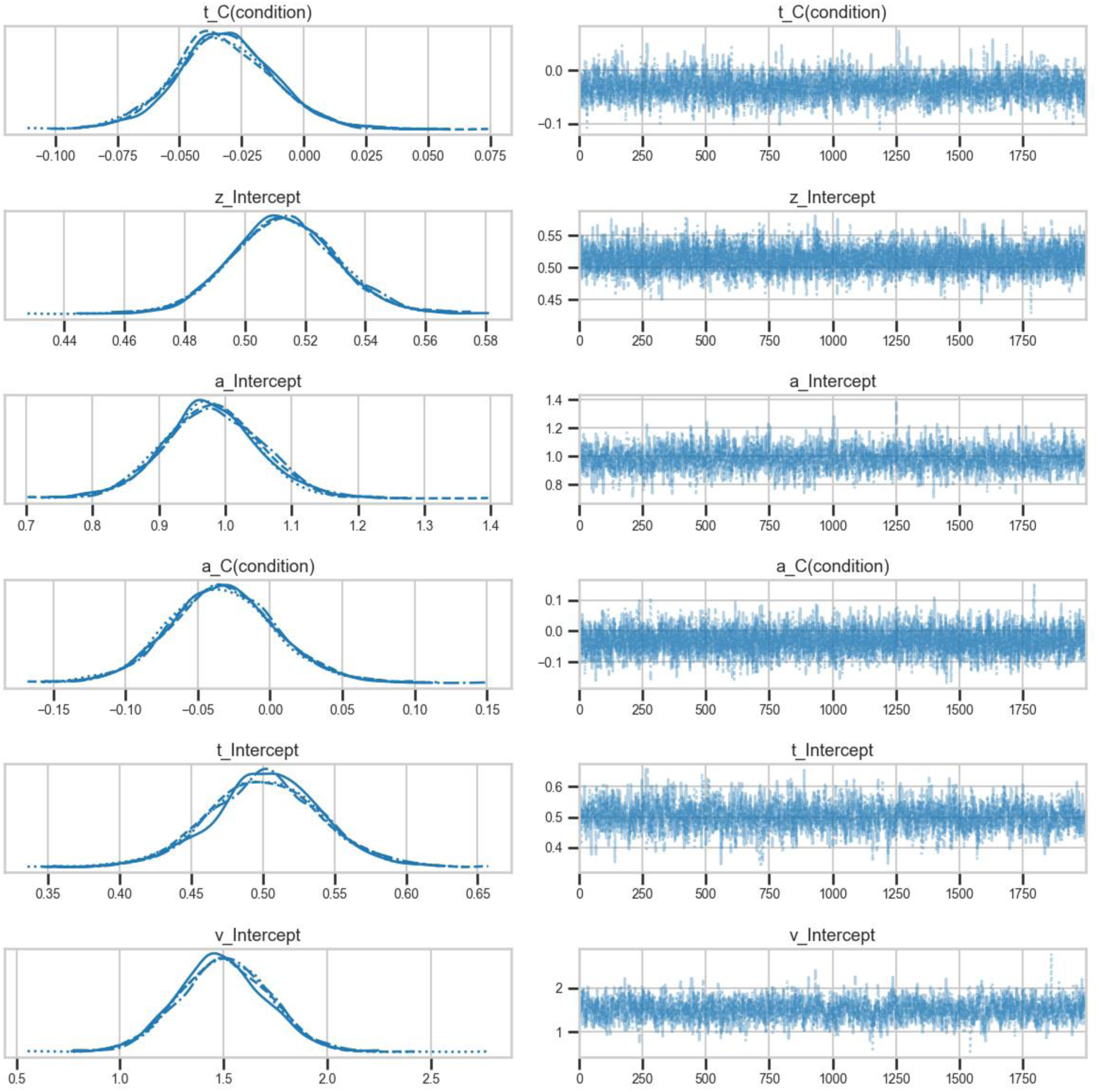
Model *at* traces

**Table S4.**
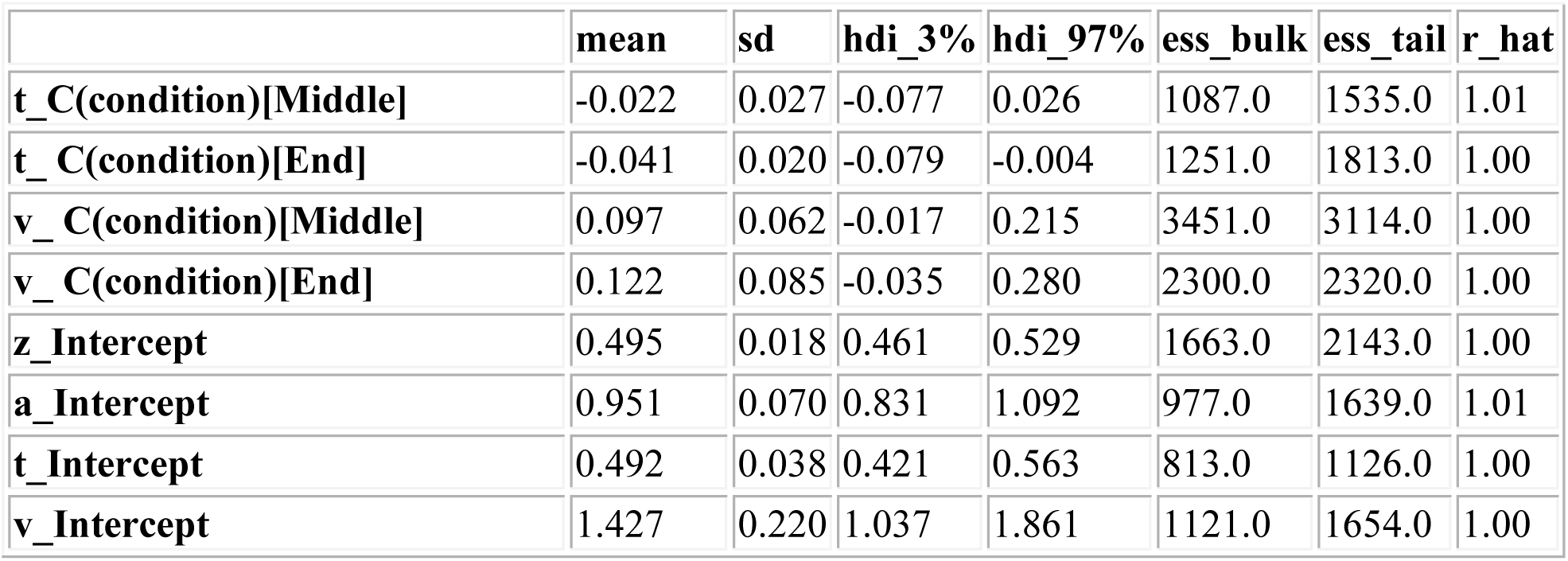
Full summary of the results of the fit of the best model (*vt* model)

**Figure S5.**
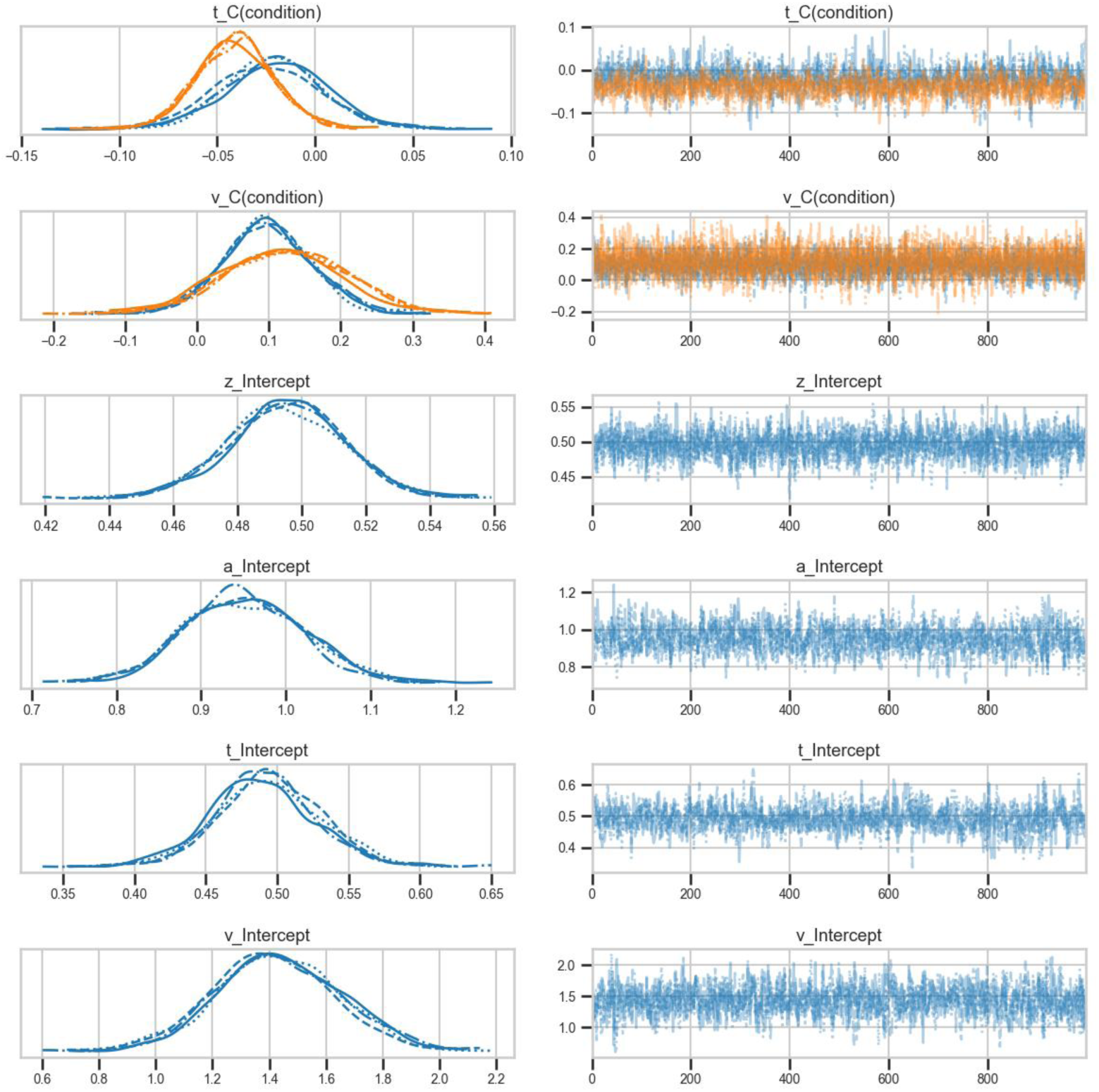
Model *vt* traces

**Figure S6.**
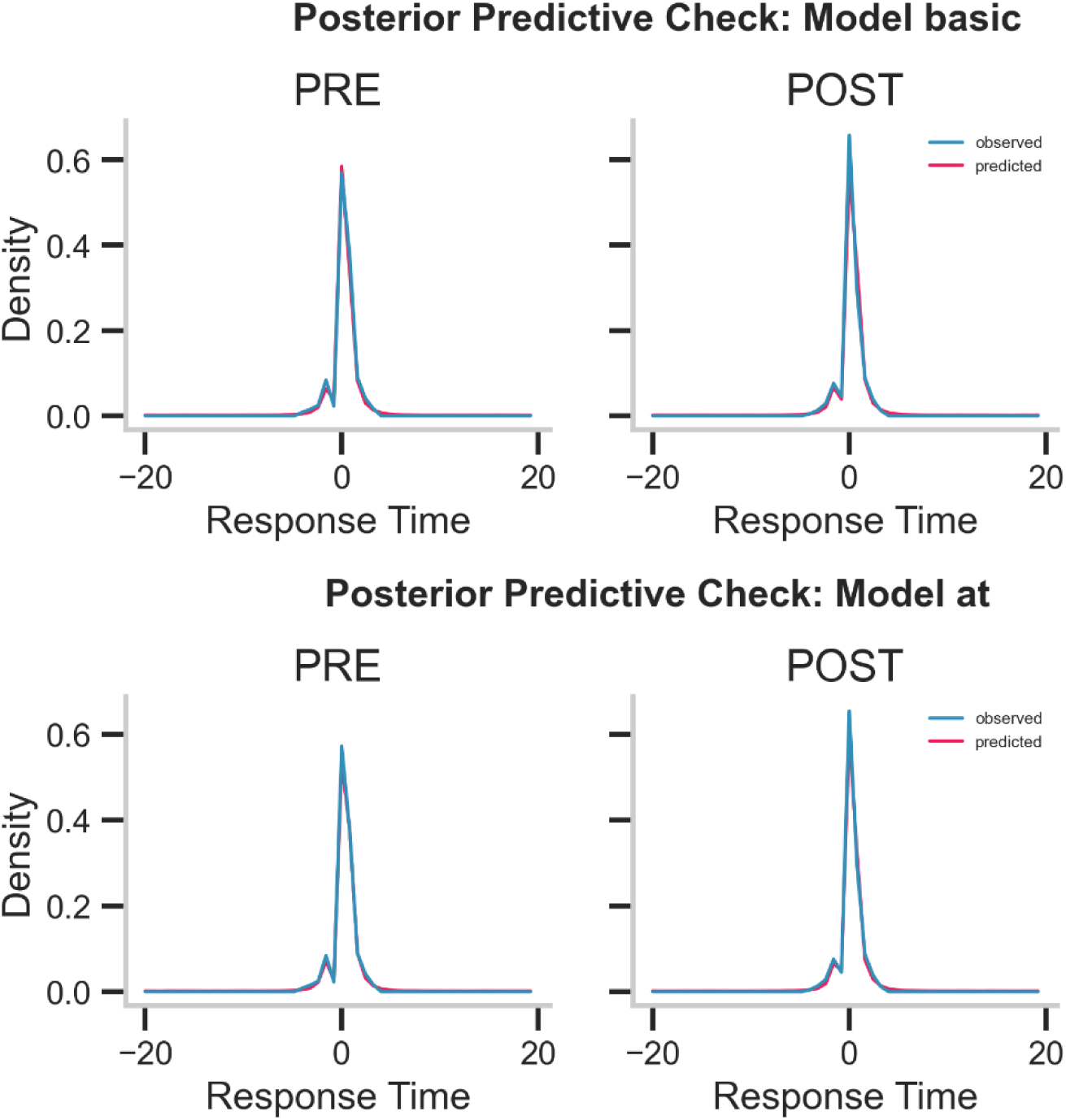
Posterior predictive check of reaction times for PRE and POST exercise.

**Figure S7.**
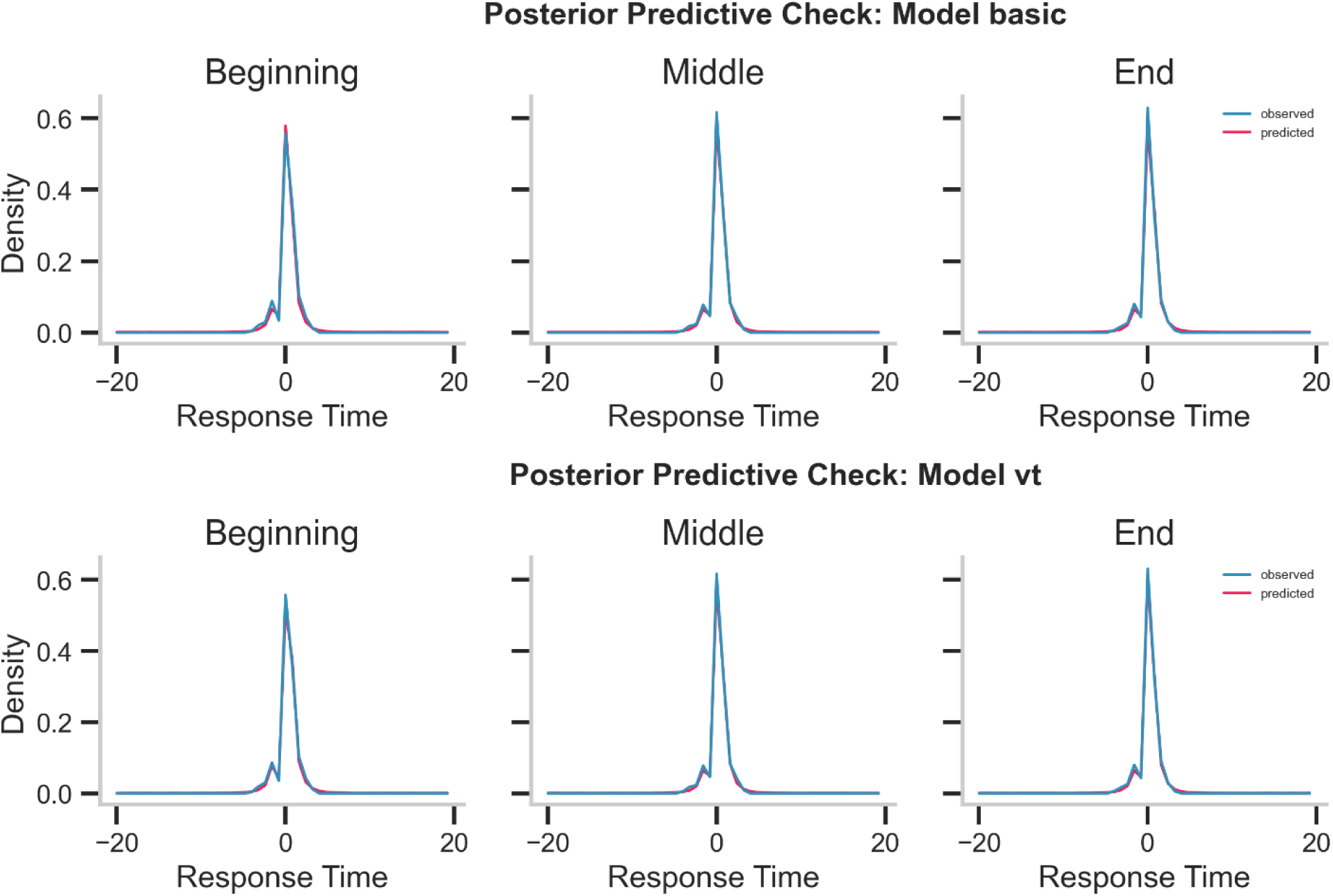
Posterior predictive check (PPC) of reaction times across exercise blocks: Beginning, Middle, and End.

## Notes

### Competing Interest Statement

The authors have declared no competing interest.

### Summary of Updates

We have revised the manuscript to clarify the RDK task and DDM parameters, replaced subjective terms with objective descriptions, addressed physiological covariates and inter-individual variability, updated figures and supplementary materials, and fully reorganized the modeling code and data repository for clarity and reproducibility (now on Zenodo: https://zenodo.org/records/17447883).

